# Genome wide identification of QTL associated with yield and yield components in two popular wheat cultivars TAM 111 and TAM 112

**DOI:** 10.1101/2020.07.27.222703

**Authors:** Yan Yang, Smit Dhakal, Chenggen Chu, Shichen Wang, Qingwu Xue, Jackie C. Rudd, Amir M.H. Ibrahim, Kirk Jessup, Jason Baker, Maria Pilar Fuentealba, Ravindra Devkota, Shannon Baker, Charles D. Johnson, Richard Metz, Shuyu Liu

## Abstract

Two drought-tolerant wheat cultivars, ‘TAM 111’ and ‘TAM 112’, have been widely grown in the Southern Great Plains of the U.S. and used as parents in many wheat breeding programs worldwide. This study aimed to reveal genetic control of yield and yield components in the two cultivars under both dryland and irrigated conditions. A mapping population containing 124 F_5:7_ recombinant inbred lines (RILs) was developed from the cross of TAM 112/TAM 111. A set of 5,948 SNPs from the wheat 90K iSelect array and double digest restriction-site associated DNA sequencing was used to construct high-density genetic maps. Data for yield and yield components were obtained from 11 environments. QTL analyses were performed based on 11 individual environments, across all environments, within and across mega-environments. Thirty-six unique consistent QTL regions were distributed on 13 chromosomes including 1A, 1B, 1D, 2A, 2D, 3D, 4B, 4D, 6A, 6B, 6D, 7B, and 7D. Ten unique QTL with pleiotropic effects were identified on four chromosomes and eight were in common with the consistent QTL. These QTL increased dry biomass grain yield by 16.3 g m^−2^, plot yield by 28.1 g m^−2^, kernels spike^−1^ by 0.7, spikes m^−2^ by 14.8, thousand kernel weight by 0.9 g with favorable alleles from either parent. TAM 112 alleles mainly increased spikes m^−2^ and thousand kernel weight while TMA 111 alleles increased kernels spike^−1^, harvest index and grain yield. The saturated genetic map and markers linked to significant QTL from this study will be very useful in developing high throughput genotyping markers for tracking the desirable haplotypes of these important yield-related traits in popular parental cultivars.

## Introduction

Wheat (*Triticum aestivum* L.) is one of the most important food crops worldwide. The significance of wheat lies on its physical and chemical properties of grains, which provide over 20% of the calories and protein requirements for human nutrition. Yield is a polygenic complex trait and the most important to breeders and farmers. However, environmental conditions and the genetic-by-environmental interactions throughout all processes of vegetative and reproductive growth and development could seriously affect yield [34]. In general, grain yield can be broken into three major components as number of spikes m^−2^ (SPM), kernels spike^−1^ (KPS), and thousand kernel weight (TKW) with each controlled by multiple genes or quantitative trait loci (QTL). Interactions among QTL and between QTL and environments also modify the expression of the QTL in different genetic backgrounds (Barton and Keightley 2002). Typically, a QTL detected in one environment but not in another might be a indication of QTL × environment interaction (QEI). However, assessing the effects of such interactions is difficult due to the unpredictable random change of environments. Goldringer et al. [8] first proposed the additive and epistatic genetic variances for agronomic traits in a doubled haploid population and demonstrated that yield and its components showed either additive or additive plus epistatic effects. Significant epistasis and QEI for yield were identified subsequently in other researches [11, 29, 36, 27, 20]. Thus, dissection of QTL effects and their interactions may facilitate better understanding of the genetic control of the complex yield traits [3].

Saturated genetic linkage maps play a crucial role in QTL identification for providing measurements of the relative effects of alleles in a mapped chromosomal region as well as selectable DNA markers for breeders to integrate the traits through marker-assisted selection (MAS) [30]. More recently, single nucleotide polymorphisms (SNPs) as the common source of genetic variation among individuals of any species and the smallest unit of genetic variation with virtually unlimited numbers (Deschamps and Campbell 2010), were used to develop high-density linkage maps and QTL identification in many crops. The availability of diverse SNP genotyping platforms, particularly genotyping-by-sequencing (GBS) based on the next-generation sequencing, were facilitated in genetic dissection, marker discovery, and genomic selection of complex traits [5,10]. However, the extensive abundance of conserved repetitive element nature of the hexaploid wheat genome (~80%) has slowed the progress in SNP discovery and detection [32]. Cavanagh et al. [4] developed 9K SNP assays and constructed the first high-density wheat consensus SNP map containing 7,504 polymorphic loci. A set of 40K out of 90K SNP assay from wheat was mapped onto chromosomes [31], thus provides a powerful resource for genome-wide dissecting traits of interests and developing new tools for efficient selection in breeding. Liu et al. [16] mapped 4k to 8k array SNPs in three wheat bi-parental mapping populations.

In this study, the highly-saturated genetic maps constructed with SNPs from 90K iSelect array and double digest restriction-site associated DNA sequencing (ddRADseq) were used to dissect QTL associated with yield, yield components, and other agronomic traits in popular cultivars TAM 111 and TAM 112. Additionally, through extensive analysis of additive-by-environment interactions, epistasis, and epistasis-by-environment interactions in individual and mega environments, the consistent and pleiotropic QTL were identified and summarized.

## Materials and Methods

### Plant Material and Phenotyping

A population of 124 F_5:7_ recombinant inbred lines (RILs) was derived from the cross between TAM 112 and TAM 111. Both the parents are hard red winter wheat (HRWW) released by Texas A&M AgriLife Research, and they are the top-ranked cultivars grown in the U.S. Great Plains. TAM 111 has the pedigree of ‘TAM 107’//TX78V3630/ ‘Centurk78’/3/TX87V1233 with excellent performance under both drought and irrigated conditions, whereas TAM 112 has the pedigree of U1254–7–9–2–1/TXGH10440 and is highly adapted to drought condition [13, 28]. Genetic analysis of the population thus can detect the favorable alleles from the two parents.

The 124 recombinant inbred line (RILs) of TAM 112/TAM 111 along with their parents were evaluated for yield and yield component traits in field experiments across 11 environments during five crop years harvested in 2011, 2012, 2013, 2014, and 2017. The combination of the location-year-irrigation level is an environment. Field locations used in this study included Texas AgriLife Research stations in Bushland (35° 06’ N, 102° 27’ W) in 2011, 2012 and 2017 (designated as 11BD, 12BD for dryland and 17BI for irrigated, respectively), Chillicothe (34° 07’ N, 99° 18’ W) in 2012 and 2014 (designated as 12CH and 14CH, respectively), two irrigation levels (75% and 100%) in Etter (35° 59’ N, 101° 59’ W), TX in 2013 and 2014 (designated as 13EP4, 13EP5, 14EP4 and 14EP5, accordingly), and Clovis (34° 24’ N, 103° 12’ W), NM (designated as 17CVI), Dumas (35° 51’ N, 101° 58’ W) (designated as 17DMS), TX in 2017. All trials were planted using alpha lattice design with an incomplete block size of five plots, and each trial has two replications in every environment. Standard agronomic practices were carried out for each environment. The data collection followed similar procedures as outlines by Assanga et al. [1]. Plot grain yield from combine harvester (YLD) was recorded. Biomass sample harvested from a random half-meter inner row showing uniform plant performance from each plot was oven-dried for 72 hrs at 60 °C and used to measure total dry biomass (BM), grain weight from the biomass sample (BMYLD), and yield components. Thousand kernel weight (TKW) was calculated by the weight of 200 seeds and scaling to 1000 seeds from biomass sample; the number of spikes m^−2^ (SPM) was calculated from the plot sample by counting the number of heads. Kernels spike^−1^ (KPS) was calculated using BMYLD, TKW and SPM; the harvest index (HI) was calculated as grain weight (BMYLD) divided by total weight of biomass sample (BM) from each plot. Single head dry weight (SHDW) was calculated through dividing the total dry head weight including glumes and awns per plot biomass sample by the number of heads. Single head grain weight (SHGW) was calculated by dividing the total BMYLD by the number of heads.

### DNA extraction and genotyping

Whole genomic DNA was extracted from leaf samples of parents and 124 RILs using the CTAB method with minor modification as described by Liu et al. [15]. SNP genotyping with Infinium iSelect assays containing 90K SNPs was performed in USDA Small Grains Genotyping Laboratory at Fargo, ND according to manufacturer’s protocol (Illumina Inc., San Diego, CA, USA), and the assay was designed under the International Wheat SNP Consortium protocols [4]. The fluorescence signal was captured by Illumina scanner and analyzed using GenomeStudio software (www.illumina.com). More details for polymorphic SNP sorting and conversion in this population were outlined in Liu et al. [16] and Dhakal et al. [6].

The 124 RILs and two parents were also genotyped in Texas AgriLife Research Genomics & Bioinformatics Services at College Station, TX (http://www.txgen.tamu.edu/) following the Double digest restriction-site associated DNA (ddRADSeq) method with some noted modifications. The libraries were constructed using a 96-plex plate with single random blank well included for quality control. DNA was co-digested with the restriction enzymes *Pst*I (CTGCAG) and *MspI* (CCGG), and barcoded adapters were ligated to individual samples. SNP calling was processed as described by Yang et al. [35].

#### Adapters

All oligos were purchased from Integrated DNA Technologies (IDT), and were received as a 100 μM stock in IDTE. Adapters were made by mixing equimolar amounts (30 μM of top and bottom oligos in 100 μl 1X annealing buffer (10mM Tris-HCl, pH 8.0, 50 mM NaCl, 1 mM EDTA). The oligos were held at 95°C for 10 sec, then cooled to 12°C at a rate of 0.1°C per sec. P5-Index Adapters were made by annealing the following oligos (where XXXXXXXX represents 8-base i5 index sequences): Top (5’ to 3’): AAT GAT ACG GCG ACC ACC GAG ATC TAC ACX XXX XXX XTC TTT CCC T, Bottom (5’ to 3’): /5Phos/AXX XXX XXX GTG TAG ATC TCG GTG GTC GCC GTA TCA TT. The P5-*Pst*I-Bridge adapters was made by annealing top (Pster_T, 5’ to 3’): /5Phos/ACA CGA CGC TCT TCC GAT CTT GCA and bottom (Pster_B, 5’ to 3’): AGA TCG GAA GAG CGT CGT GTA GGG AAA G oligos. *P7-Mlu*CI Adapter was made by annealing top (P7-*Mlu*CI_T, 5’ to 3’): AAT TAG ATC GGA AGA GCA CAC GTC TGA ACT CCA GTC AC and bottom (P7-MluCI_B, 5’ to 3’): GTG ACT GGA GTT CAG ACG TGT GCT CTT CCG ATC T.

#### Dual Ligation ddRAD

At the end of each step in this protocol, samples were quantified on a DeNovix DS-11 spectrophotometer. One hundred nanograms of DNA per sample in 96 well plate format was digested in a final volume of 25 μl in 1X NEB Cut Smart Buffer and 200 U *Pst*I-HF and 100 U *Mlu*CI (New England Biolabs) at 37°C for 4 hours. Following a 20 min 80°C enzyme inactivation, samples were held at 12°C until ligation. To each 25 μl digest, 5.7 μl of a master mix was added such that each well got the equivalent of 3.2 μl 10X Ligase buffer (NEB), 0.25 μl T4 DNA Ligase (New England Biolabs) and P5-*Pst*I-Bridge and *P7-Mlu*CI adapters at a final concentration of 500 nM. In addition, each well got 1 of 48 unique P5-Index Adapters (400nM final concentration) and were mixed by pipetting. Plates were spun down and incubated at 16°C for 8 hr followed by a 15 min heat inactivation at 65°C and held at 4°C. Pools were made by combining no more than 48 samples with unique P5 indexes. To each pool, EDTA was added to 0.25mM to further inhibit ligase activity. Pools were precipitated by adding 1/10^th^ volume of 3M sodium acetate (pH 5.2), evenly dividing them into two or three 2.0 ml microcentrifuge tubes and adding 2 volumes 100% ethanol and placing them at −20°C for at least 1 hr. Tubes were spun at 20,000 xg for 10 min and supernatant poured out. Tubes were washed with 1 ml 70% ethanol, spun 5 min and supernatant removed. Pellets were resuspended in 200μl EB and purified through QIAquick PCR Purification Kits as per manufacturer’s protocol (Qiagen) eluting twice with 50 μl EB. Combined elutants (100 μl total) were further purified to remove unligated adapters using 0.9X volume AMPure XP beads as per the manufacturer’s protocol (Beckman-Coulter) eluting in 35μl EB.

Up to 3000ng of each pool was size selected at 390-610 bp (280-600 bp inserts plus 110bp adapters) on Pippin prep 2% dye-free gels (Sage Science). Recovered DNA was purified with 0.9X AMPure XP beads as described earlier (Beckman-Coulter) and eluted in 32μl EB.

Incorporation of a biotin moiety at the P5 side (for further purification - described later) and the addition of the i7 index was accomplished in a Pre-Selection PCR step. Using the primers P5_Select (5’-3’): /5BiotinTEG/AAT GAT ACG GCG ACC ACC GAG ATC TAC AC and one of twenty four i7 indexed reverse primers (TDX 1-24, 5’-3’ where XXXXXXX represent bases used for indexes: CAAGCA GAA GAC GGC ATA CGA GAT XXX XXX XGTGAC TGG AGT TCA G AC GTG TGC). PreSelect PCR reactions (200 μl total volume, split into two reactions of 100 μl each) contained up to 150 ng size-selected DNA, 0.4 mM dNTPs, 1 μM each primer (P5-Select and a TDX-reverse index), 20 U of Q5 Polymerase in 1X Q5 High Fidelty DNA Polymerase Buffer. Reactions were denatured at 98°C for 30 sec, then subjected to 15 cycles of 98°C for 10 sec, 62°C for 20 sec and 72°C for 40 sec with a final elongation at 72°C for 5 min followed by a 10°C hold. Pre-Selection PCR reactions were cleaned up with QIA quick columns and AMPure XP beads as described above with a final elution in 50μl EB.

Selection of only *Pst*I-*Mlu*CI fragments was accomplished using Dynabeads M-280 Streptavidin beads (Invitrogen) to capture fragments with biotin incorporated at their P5 ends during PreSelect PCR. Dynabeads (50 μl per pool) were captured on a magnet and washed twice in 300 μl 1X bead washing buffer (10mM Tris, pH 7.5, 2 M NaCl, 1 mM EDTA) by resuspending beads in buffer, capturing on a magnet, and removing the supernatant with pipette tips. After the second wash, beads were resuspended in 100 μl 2X bead wash buffer per pool, and 100 μl washed beads was mixed with up to 3000 ng of Pre-Selected DNA (in 100 μl EB). Samples were incubated at RT for 20 min then captured on a magnet. *Mlu*CI-*Mlu*CI fragments lacking biotin were washed away as follows: beads were washed three times in 200 μl 1X bead washing buffer, twice in 200 μl nuclease free water and once in 100μl 1x SSC (150 mM NaCl, 15mM sodium citrate, pH 7.0, BIO-RAD Laboratories, Inc.) and finally resuspended in 50 μl 1X SSC. *Pst*I-*Mlu*CI fragments were obtained by heating beads at 95°C for 5 min thus denaturing off the non-biotinylated strand (leaving both strands of *Pst*I-*Pst*I fragments attached to the beads). Following 5 min at 95°C, tubes were transferred immediately to a magnet and supernatant removed quickly to new tubes. This was repeated for a total of two heated elutions totaling 100μl. Elutants were purified with QIA quick PCR columns as described above, eluted in 40 μl EB and quantified.

Final libraries were produced in a PCR reaction of 50 μl containing 10 ng single-stranded, Dynabead-selected DNA, 0.4μM dNTPS, 0.5μM each final PCR primers (DuLig-F1, 5’-AAT GAT ACG GCG ACC ACC GAG ATC TAC AC-3’ and DuLig-R1, 5’-CAA GCA GAA GAC GGC ATA CGA GAT-3’) and 20U/μl Q5 DNA Polymerase in 1X Q5 DNA polymerase reaction buffer (New England Biolabs). Reaction conditions were the same as Pre-Selection PCR, but total cycle number was 8. Final PCR reactions were cleaned up with 0.9X AMPure XP beads, eluted in 35 μl EB, quantified and assessed for quality on a Fragment Analyzer (Agilent Technologies) diluted and quantified by qPCR (Kappa Biosystems). Libraries were sequenced on the Illumina NovaSeq 6000 system, S4XP flowcell running 2X 150 bp recipe.

### Statistical analysis

The analysis of variance (ANOVA) from individual and across environment data was calculated to determine the significance of genetic (G), environment (E), and genetic-by-environment interaction (GEI) variances. Broad-sense heritability was calculated, and only single environment with heritability ≥ 0.05 were included into the analysis. Pearson’s correlation coefficients among all variables were calculated. Best linear unbiased predictors (BLUP) and best linear unbiased estimator (BLUE) of individual environment and across all environments were computed using a restricted maximum likelihood (REML) approach based on META-R program with lme4 package in R software from Matthews and Foulk [18]. Mega-environments for each trait were classified according to the biplot clustering for the environments. In most case, the BLUP values were used for the QTL analyses. The BLUE values were used only if the BLUP values were the same for all the RILs in that environment.

### Linkage map construction and QTL identification

Linkage map construction in this population has been described in Yang et al. [35]. Of the marker data generated, the false double crossovers were manually checked and removed according to the alignment of SNP orders between genetic maps and physical base pair location from IWGSC RefSeq v1.0 (https://wheat-urgi.versailles.inra.fr/Seq-Repository/Assemblies, accessed on February 8, 2020). QTL analysis was performed using QTL IciMapping software [19]. Individual environment QTL analysis was conducted using single trait from single individual environment and across multiple environments. The multi-environment QTL analyses were also performed using single trait across classified mega-environments and within each mega-environment. The genetic position and effects of individual environment QTL and multi-environment trait QTL were determined by integrated composite interval mapping (ICIM) function for additive effect (ICIM-ADD) and epistasis effect (ICIM-EPI). To identify an appropriate threshold likelihood of odd (LOD) score for declaring a significant QTL, permutation test was conducted for 1,000 times for ICIM-ADD for individual and across environments. Consistent QTL was determined if a QTL was significant at least from two individual environments or two out of the four analyses including individual environment, across all individual, within and across mega environments. Pleiotropic QTL was determined if a QTL was significantly associated with two or more traits that were not highly correlated to each other. For ICIM-EPI, since it is too long to run permutation, LOD = 5 was chosen but actual thresholds for each trait from ICIM-ADD as reference or 10 if too many interactions exist.

Identified QTL were designated in the format as *Qtrait.tamu.chrom.Mb,* where *trait* is an abbreviation for a trait name, *tamu* represents Texas A&M University, *chrom* is chromosome on which the QTL is located, and *Mb* is a physical position of the QTL peak linked marker according to alignment with the IWGSC RefSeq v1.0 reference genome (International Wheat Genome Sequencing Consortium, 2014).

## Results

### Trait analysis

The combined ANOVA across environments indicated a significant genetic variance for all traits except dry biomass and significant genetic by environment interactions (GEI) except spikes m^−2^ (*P* < 0.01) (Table S1).

The entry-mean heritability ranged from moderate (0.4 to 0.6) to high (> 0.6). Yield and three yield components, including thousand kernel weight, kernels spike^−1^, spikes m^−2^, as well as single head dry weight and single head grain weight, exhibited higher heritability (0.75-0.90). Harvest index displayed moderate heritability (0.56), whereas dry biomass and biomass grain yield expressed relatively low heritability of 0.23-0.33. For the overall BLUP means of three duplicated sets across seven to 11 environments, TAM 112 had higher dry biomass and spikes m^−2^ while TAM 111 had higher kernels spike^−1^, single head dry weight and grain weight (Table S1).

Based on the best linear unbiased prediction (BLUP) means, positive genetic correlations were found between yield and the three yield components expect thousand kernel weight (TKW) (Table S2). Dry biomass had significant positive correlations with all tested traits except thousand kernel weight and spikes m^−2^. Harvest index had significant correlations with all traits except spike m^−2^. Thousand kernel weight only had a significant low positive correlation with harvest index but a significant negative correlation with spikes m^−2^. Spikes m^−2^ had a significant negative correlation with kernels spike^−1^. Thousand kernel weight had the least significant correlations with other traits related to yield indicated that it could be improved independently. However, kernels spike^−1^ can be increased along with thousand kernel weight for improved yield but not together with spikes m^−2^ due to the significant negative correlations. Across all individual 11 environments, harvest index were significantly and positively correlated with all the traits except thousand kernel weight in 12BD, 17CVI, and 17BI (significantly negative), spikes m^−2^ in 12BD, 12CH, 17DMS, 11BD and 13EP5 (last two significantly negative), kernels spike^−1^ in 14CH, and yield in 13EP5 (Table S2); thousand kernel weight was significantly and negatively correlated with most of the rest traits except kernels spike^−1^ in 12BD, 12CH, 13EP5, 17CVI, 17DMS, 13EP4, 14CH, and 14EP5 (last three significantly positive), spikes m^−2^ in 14CH and 14EP5, yield in 13EP4, 14EP4, 17CVI, 12BD, 12CH, 14CH 14EP5, and 17DMS (last five significantly positive); spikes m^−2^ was significantly and negatively correlated with kernels spike^−1^ in 11BD,12BD, 12CH, and 13EP4, except in 14CH, 17DMS, and 13EP5 (last one significantly positive), significantly and positively correlated with yield in 11BD, 12BD, and 14CH, except in 12CH, 13EP4, 13EP5, and 14EP5 (last two significantly negative); kernels spike^−1^ was significantly and positively correlated with yield in all 11 environments (Table S2).

The significant correlations between yield and its components implied that yield can be increased through the indirect selection of the higher component traits. Hence, mapping the QTL for yield and associated yield components could reveal significant QTL across environments and improve the indirect selections.

### Boxplot and biplot across all environments, and mega environment classification

From the boxplot of all the traits across individual environments, it is easy to define that the lower yield environments were from the drought years (eight environments from 11BD to 14EP5) while the higher yield environments were from the irrigated location in a good rainfall year (three environments, 17BI, 17CVI, and 17DMS) with the latter had a higher genetic variations (Table S1a). Similar trends were found on dry biomass and biomass grain yield; however, several other traits did not follow this trend. The three environments having higher harvest index were 11BD, 14EP4, and 14EP5, and all the environments had relatively larger variations, ranging from 20% to 50%. Kernels spike^−1^ had very similar means across all the environments except 17BI and 17DMS. Thousand kernel weight were classified into two groups and the higher median group included all the irrigated environments in 2014 and 2017. However, spikes m^−2^ did not have similar trends as any other traits. Its median and ranges were very similar in the two irrigated environments, 17BI and 17DMS (Fig S1a).

Biplot of all the environments for each trait could help us to classify those environments where the performance of individual lines had similar trends; therefore, we classified them as a mega environment (ME) (Fig S1b). Yield had ME1 (17BI, 17CVI, 14EP4, 14EP5); ME2 (12BD, 13EP4, 13EP5), and ME3 (12CH, 14CH); dry biomass had ME1 (11BD, 14CH, 14EP4) and ME2 (13EP4, 13EP5); biomass grain yield had ME1 (11BD, 14CH), ME2 (13EP4, 12BD, 12CH), ME3 (14EP4, 14EP5, 17CVI); harvest index had ME1 (14EP5, 17BI, 17CVI, 17DMS), ME2 (12CH, 14CH, 13EP4), ME3 (12BD, 11BD, 13EP5); kernels spike^−1^ had ME1 (17CVI, 17DMS, 14EP4, 14EP5), ME2 (11BD, 12BD, 13EP4, 14CH); spikes m^−2^ had ME1 (14EP5, 14CH), ME2 (12BD, 13EP4, 13EP5, 17CVI), ME3 (12CH, 17DMS); thousand kernel weight had ME1 (11BD, 12BD, 12CH, 13EP4, 13EP5, 14CH), ME2 (14EP4, 14EP5, 17BI, 17CVI, 17DMS). The mega environments allowed us to identify some consistent genetic factors across similar individual environments, within and across mega environments.

### Linkage map

A set of 5,948 markers including 3,193 from ddRADseq and 2,740 from 90K iSelect array SNPs, and 15 microsatellites and kompetitive allele specific PCR (KASP) markers were used to construct 25 linkage groups covering all 21 chromosomes were used for QTL analyses (Table S3). The cumulative genetic map length is 2,703.9 cM with an average marker density of 0.6 SNP/cM or 2.8 SNP/Mb. The total covered physical base pair length is about 12.6 Gb with average length of 602.2 Mb per chromosome.

### Consistent QTL identification for individual trait

A set of 87 unique QTL regions significantly associated with nine yield and yield related traits across 11 environments over five years were identified through the analyses of data from individual and mega-environments (Table S4; Fig S2 and S3). Among them, a set of 36 unique consistent QTL was identified to be associated with one trait but from at least two out of the analyses from individual, across all individual, within and across each defined mega environments based on biplot and overall best linear unbiased prediction (BLUP) or best linear unbiased estimation (BLUE) for each trait (Tables 1 and 2; Fig 1). A set of 10 unique pleiotropic QTL was found to be associated with at least two traits that were not highly correlated to each other (Table 2). Among the consistent and pleiotropic QTL, eight were in common (Tables 1 and 2; Fig 1).

**Fig 1.**
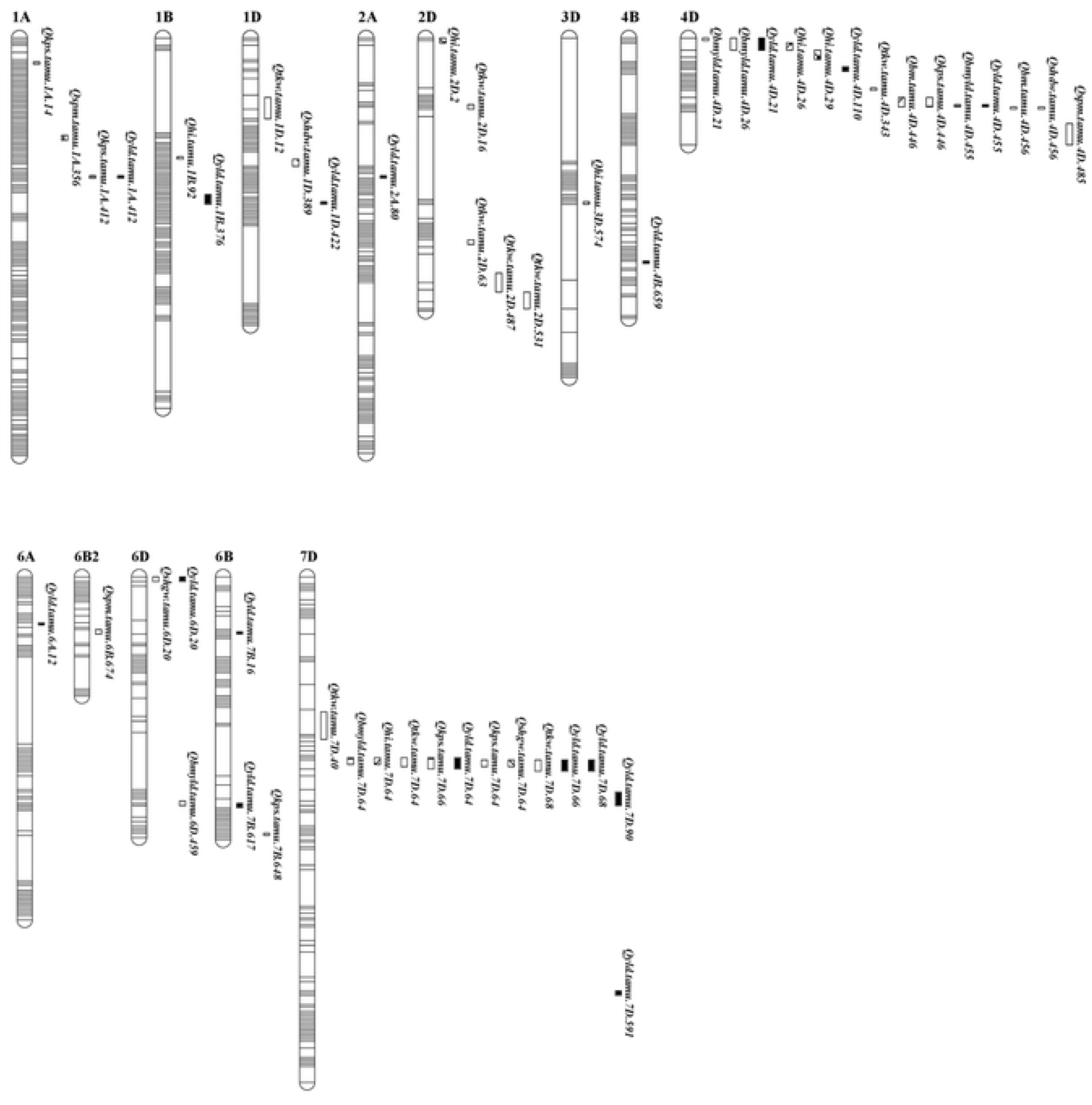
Consistent and pleiotropic QTL identified from individual and mega-environinents for all traits. Traits include 1) Yield from combine plots (YLD), 2) dry biomass from hand harvested 0.5 m long inner row sample from crown (BM), 3) grain weight from b) as hand harvested dry grain (BMYLD), 4) harvest index (HI), 5) kernels spike’^1^ (KPS), 6) spikes nr^2^ (SPM), 7) thousand kernel weight (TKW), 8) single head dry weight (SHDW), 9) single head grain weight (SHGW). Identified QTL were designated in the format as *Otrait, tamu.chrom.Mh.*

**Table 1.**
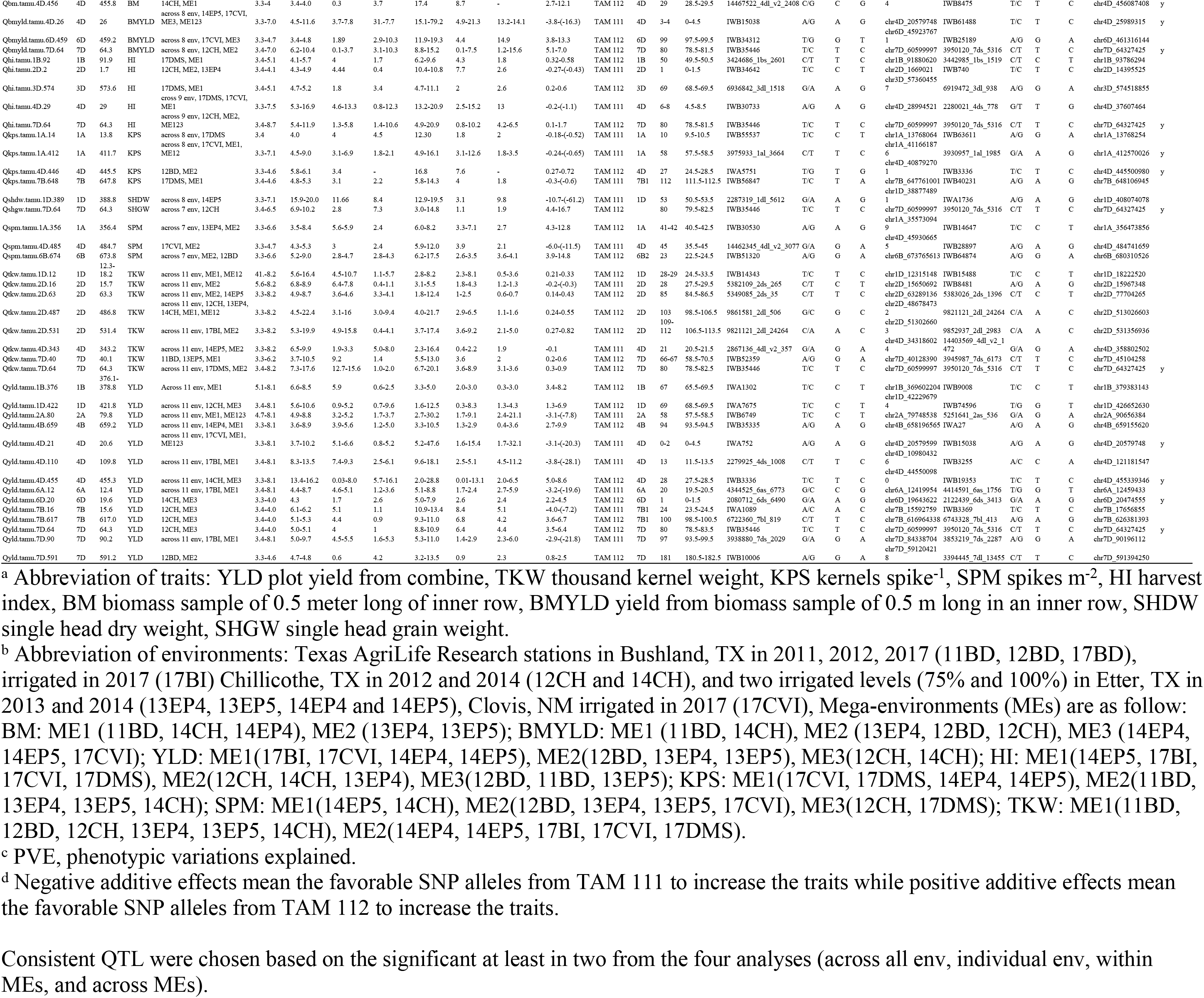
Consistent QTL associated with yield and yield components from at least two analyzed across individual or mega environments in TAM 112/TAM 111

**Table 2.**
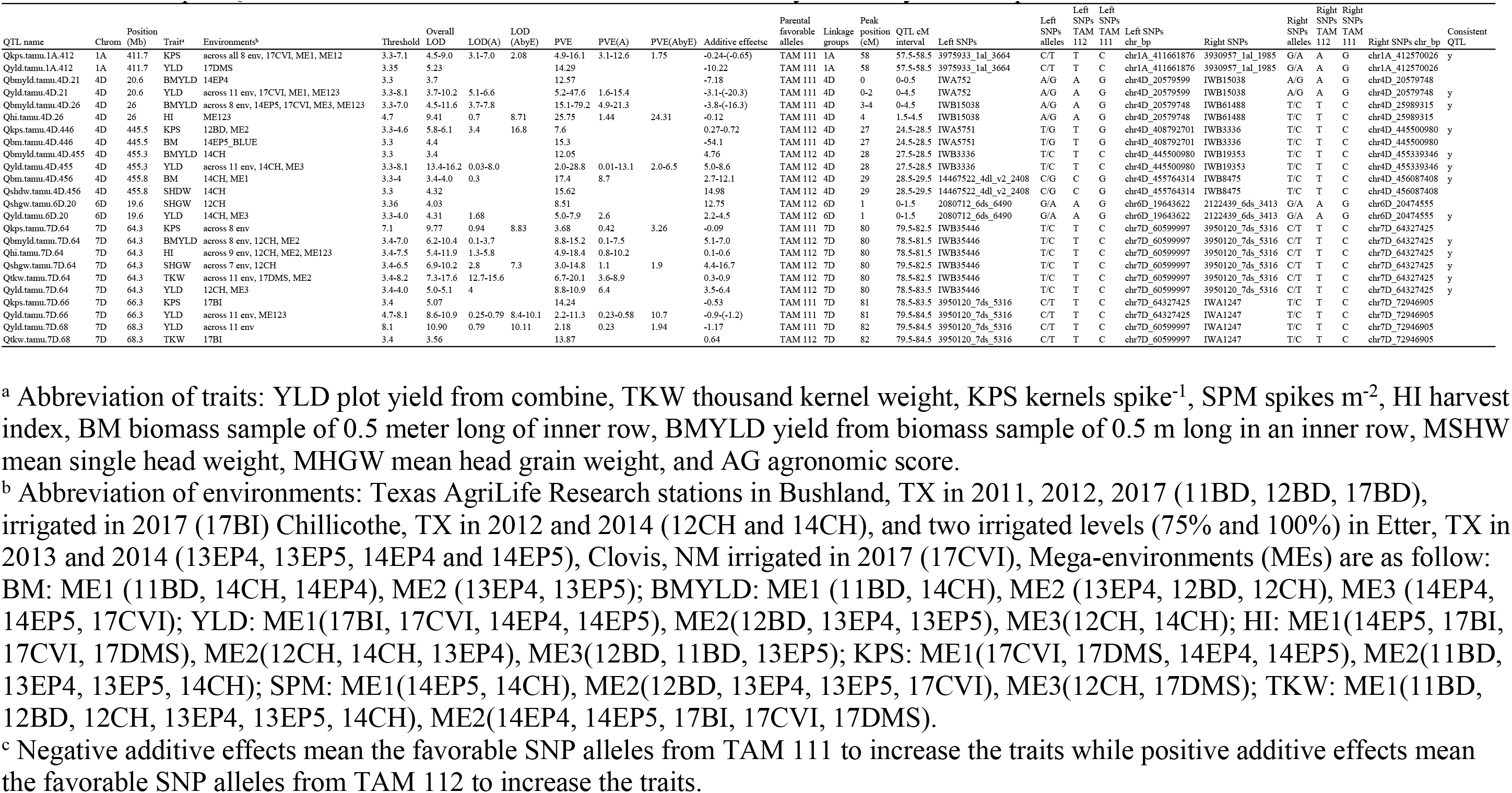
Pleiotropic QTL associated with at least two different traits of yield and yield components in TAM 112/TAM 111

### Yield

A set of 14 consistent QTL for yield was identified on chromosomes 1B, 1D, 2A, 4B, 4D, 6A, 6D, 7B and 7D (Table 1, Table S4). There were four major QTL at 20.6 and 109.8 Mb on 4D, 12.4 Mb on 6A, and 90.2 Mb on 7D that increased yield up to 19.6 - 28.1 g m^−2^ from the analyses of individual environment 17BI or 17CVI, ME1 (including 17BI, 17CVI, 14EP4, and 14EP5) and across 11 environments and all had favorable alleles from TAM 111. Eight minor QTL with favorable alleles from TAM 112 that increased yield by 2.5 - 9.9 g m^−2^ were located at 376.1 Mb on 1B, 421.8 Mb on 1D, 659.2 Mb on 4B, 455.3 Mb on 4D, 19.6 Mb on 6D, 617.0 Mb on 7B, and 64.3 Mb and 591.2 Mb on 7D. From the LOD score and R^2^ values of additive effects, only five out of the eight minor QTL, *Qyld.tamu.1B.376, Qyld.tamu.1D.422, Qyld.tamu.7B.16, Qyld.tamu.7B.617,* and *Qyld.tamu.7D.64* had larger proportion of additive effects while the rest had larger additive-by-environment interactions than additive effects indicating the complex of yield inheritance. Among the four major QTL that had larger additive effects, results from across individual environment analyses showed that the corresponding additive-by-environment interactions increased yield by 15.8 - 24.0 g m^−2^ at 17BI or 17CVI (Table S4).

### TKW

Eight QTL were identified for thousand kernel weight including one on chromosome 1D at 12.3 Mb, four on 2D at 15.7, 63.3, 486.8 and 531.4 Mb, one on 4D at 343.2 Mb, and two on 7D at 40.1 and 64.3 Mb (Table 1). Two QTL *Qtkw.tamu.2D.16* and *Qtkw.tamu.4D.343* had the favorable alleles from TAM 111 and increased TKW up to 0.3 g while the other six QTL had alleles from TAM 112 and increased TKW up to 0.9 g. All QTL appeared across 11 environments and ME2 (including 14EP4, 14EP5, 17BI, 17CVI, and17DMS) analyses except three QTL *Qtkw.tamu.1D.12, Qtkw.tamu.2D.487,* and *Qtkw.tamu.7D.40.* Four major QTL *Qtkw.tamu.2D.487, Qtkw.tamu.2D.531, Qtkw.tamu.7D.40* and *Qtkw.tamu.7D.64* increased thousand kernel weight from 0.6 to 0.9 g at 12CH, 17CVI, 11BD, and 17DMS, respectively (Table S4). Their corresponding additive-by-environment interactions increased thousand kernel weight by 0.6, 0.3, 0.5, and 0.3 g, respectively (Table S4).

### KPS

Only four QTL significantly associated with kernels spike^−1^ were identified on chromosomes 1A at 13.8 Mb and 411.7 Mb, 4D at 445.5 Mb, and 7B at 647.8 Mb (Table 1). All favorable alleles were from TAM 111 and increased kernels spike^−1^ up to 0.7 except the *Qkps.tamu.4D.446.* Two of the three QTL appeared in the analyses of across eight environments, or either 17CVI or 17DMS and ME1 (including 17CVI, 17DMS, 14EP4, and 14EP5). The corresponding additive-by-environment interactions of the three QTL increased kernels spike^−1^ by 0.3 - 0.4 at 17DMS or 17CVI (Table S4).

### SPM

For spikes m^−2^, three QTL were detected on chromosomes 1A at 356.4 Mb, 4D at 484.7 Mb, and 6B at 673.8 Mb (Table 1). *Qspm.tamu.1A.356* and *Qspm.tamu.6B.674* had alleles from TAM 112 and increased spikes m^−2^ by 14.8 while *Qspm.tamu.4D.459* had allele from TAM 111 and increased spikes m^−2^ by 11.5 at 17CVI. All three QTL appeared in the analyses of ME2 (including 12BD, 13EP4, 13EP5, 17CVI) and the two QTL on 1A and 6B appeared in the analyses of across seven environments. *Qspm.tamu.6B.674* had the highest additive effects of 14.8 from 12BD and its additive-by-environment interactions increased spikes m^−2^ by 10.6 while *Qspm.tamu.1A.356* increased spikes m^−2^ by 12.8 and its interactions at 13EP4 increased 8.3 (Table S4).

### HI

Five QTL for harvest index were detected, in which the two QTL at 1.7 Mb on chromosome 2D and at 29 Mb on 4D had favorable alleles from TAM 111. *Qhi.tamu.4D.29* increased harvest index by 1.1% and were consistent in two individual environments, 17CVI and 17DMS. The other three QTL on chromosomes 1B, 3D and 7D had favorable alleles from TAM 112 and increased harvest index by 0.6% at environments 17DMS or 12CH. These five QTL could increase harvest index by 0.25 to 0.87 from additive-by-environment interactions at their corresponding environments, 17DMS, 17CVI or 12CH (Tables 1 and S4).

### BMYLD

For biomass grain yield (BMYLD) collected from 0.5-m long in an inner row, three significant QTL were identified on chromosomes 4D, 6D, and 7D (Table 1). Qbmyld.tamu.4D.26 had favorable allele from TAM 111 and increased biomass yield by 16.3 g m^−2^ at two individual environments 17CVI and 14EP5. The other two QTL at 459.2 Mb on 6D and 64.3 Mb on 7D had favorable alleles from TAM 112 and increased biomass yield up to 13.3 g m^−2^ at 12CH. Only the major QTL Qbmyld.tamu.4D.26 had a larger additive LOD scores compared with those of corresponding additive-by-environment interactions that increased biomass yield by 7.6 and 11.9 g m^−2^ at environments 17BI and 17CVI, respectively (Table S4). On the other hand, the QTL Qbmyld.tamu.7D.64 had additive-by-environment interaction effect of 16.1 g m^−2^ with allele from TAM 111 at 17BI from the analyses of across eight environments (Table S4). At another environment 12CH, the additive-by-environment interaction of the same QTL increased 7.7 g m^−2^ with allele from TAM 112. Only one significant QTL for dry biomass at 455.8 Mb on chromosome 4D and it increased biomass by 12.1 g m^−2^ with favorable allele from TAM 112 (Table 1).

In general, we can see that TAM 111 favorable alleles mainly increased kernels spike^−1^ while TAM 112 favorable alleles mainly increased spikes m^−2^ and thousand kernel weight. For biomass yield, yield, and harvest index, almost half QTL had TAM 111 favorable alleles and half had TAM 112 alleles (Table 1). A major QTL had the highest additive effects for certain trait at a particular environment. In the meantime, it had a higher effect from additive-by-environment interactions at the same environment.

### Pleiotropic QTL

A set of ten unique significant QTL regions was found to affect more than one trait and thus considered having pleiotropic effects (Table 2 and Fig 1). Eight were in common with the 36 consistent QTL identified for all nine evaluated traits. They were the QTL at 411.7 Mb on 1A that was linked to yield and kernels spike^−1^ with all the favorable alleles increasing the traits from TAM 111; the QTL at 20.6 Mb on 2D that was linked to both yield and biomass yield and the QTL at 26 Mb on 4D that was linked to yield and harvest index with all favorable alleles from TAM 111; two additional QTL at 455.3 and 455.8 Mb on 4D that were linked to biomass grain yield and yield, dry biomass and single head dry weight, respectively, with all favorable alleles from TAM 112; the fifth QTL on 4D at 445.5 Mb that increased kernels spike^−1^ by 0.72 with favorable allele from TAM 112 while it increased dry biomass by 54.1 g m^−2^ based on the BLUE value from 14EP4 (Table 2 and Table S4); the QTL at 19.6 Mb on 6D that was associated with yield and single head grain weight with favorable alleles from TAM 112; the QTL at 64.3, 66.3, and 68.3 Mb on 7D that were linked to yield, biomass yield, thousand kernel weight, kernels spike^−1^, and harvest index with the most favorable alleles increased yield and kernels spike^−1^ from TAM 111 while the favorable alleles increased thousand kernel weight and harvest index from TAM 112 (Table 2). The last two were not consistent QTL (Tables 1 and 2).

### Epistasis, epistasis-by-environment, and additive-by-environment interactions

Only those with overall LOD scores > 5.0 were summarized for the epistasis and additive-by-environment interactions (Table S5). Among 375 interactions for yield, only 56 had overall LOD scores >= 10.0, but none of the epistasis and additive-by-environment interactions had LOD > 10.0 (Fig S4). Among 28 interactions that increased yield by more than 10 g m^−2^, there were six additive-by-environment interactions at 17BI with favorable alleles from TAM 111 that increased yield from 10.4 to 17.5 g m^−2^; two additional additive-by-environment interactions at 14EP4 and 17CVI, respectively increased yield by 10.2 and 10.4 g m^−2^ with favorable alleles from TAM 112. Among 19 epistasis-by-environment interactions, 17 interactions at 17BI increased yield by 10.1 and 13.1 g m^−2^ with seven favorable alleles from TAM 112 and 10 favorable alleles form TAM 111.

Among 234 interactions for spikes m^−2^ with LOD > 5.0, only eight interactions had overall LOD score >= 10.0 and no epistasis LOD >= 10.0. Five epistasis-by-environment interactions occurred in ME2 (including12BD, 13EP4, 13EP5, and 17CVI) in which three of them had favorable alleles from TAM 112 and increased spikes m^−2^ by 10.8 and two increased spikes m^−2^ by 14.4 with alleles from TAM 111 (Table S5 and Fig S4).

For thousand kernel weight (TKW), among 581 interactions with overall LOD >= 5.0, there were 123 with overall LOD > 10.0 and 26 with epistasis LOD >= 10.0. There were five epistasis that increased TKW by 0.4-0.7 g with four having favorable alleles from TAM 112. However, only two of the five had epistasis LOD >= 10.0. Two of the five occurred within mega-environment ME2 including 14EP4, 14EP5, 17BI, 17CVI, and 17DMS. Among the eight additional interactions that increased TKW by 0.4 to 0.7 g, four were epistasis-by-environment interactions with all favorable alleles from TAM 112 and three of the four occurred in 17BI while the four additive-by-environment interactions occurred in 17CVI and 11BD with three having favorable alleles from TAM 111 (Table S5 and Fig S4).

Among 243 interactions with LOD > 5.0 for kernels spike^−1^, only four had overall LOD >= 10.0 but none of them could increase the trait by > 0.4. Among six interactions that increased kernels spike^−1^ by 0.4, four epistasis-by-environment interactions increased kernels spike^−1^ by 0.4-1.0 with two having favorable alleles from TAM 111. The one increased by 1.0 had favorable alleles from TAM 111 at 13EP5 while the same interaction increased 0.4 with favorable allele from TAM 112 in 17DMS (Table 5 and Fig S4).

For harvest index, there were 240 interactions that had LOD > 5.0 but only one interaction had overall LOD > 10.0. Four additive-by-environment and six epistasis-by-environment interactions increased harvest index by 0.5-0.8% at 17CVI with eight having favorable alleles from TAM 111.

For biomass grain yield, 190 interactions had LOD > 5.0 but only one had LOD > 10.0. All 16 epistasis-by-environment interactions at 17BI increased biomass grain yield by 15.1 to 19.6 g m^−2^ with ten having favorable alleles from TAM 112 and six having favorable alleles from TAM 111 (Table S5).

For total dry biomass, no interaction had LOD >= 10.0. Only four interactions explained 10.3 - 12.2% of total phenotypic variations but none of epistasis and epistasis-by-environment interactions increased more than 10 g m^−2^ (Table S5).

## Discussion

### Evaluation of yield and yield component in individual and mega-environments

Yield is a complex trait affected by genetic, environment and genetic-by-environment interactions. Management in crop growing conditions, such as drought or irrigated, can also interfere with grain yield. Therefore, yield trials from multiple years at multiple locations are crucial to provide data of yield and yield components under various weather and management conditions including dryland and irrigating, and further lead to more reliable genetic analysis for yield plasticity [9]. In this study, we used an alpha lattice experimental design to conduct the trials in five growing seasons and up to five locations, which provided diverse growing conditions to evaluate yield and yield-related traits and thus being able to detect effects due to genetic and genetic-by-environment interactions. Through combined ANOVA and heritability analyses, trait data showed genetic variance at a significance level with heritability higher than 0.05 were used for QTL analysis. In addition, Pearson’s product moment correlation was conducted among all traits, and most of the correlations are significant, which was further supported by co-localized QTL linked to yield and yield components to indicate the presence of pleiotropy in genomic regions modulating the quantitative traits [17], and the positive correlation thus suggests a possible linkage existing in coupling phase or presence of positive pleiotropic effects [17].

Mega-environments (MEs) have been initially defined by CIMMYT as similar biotic and abiotic stress, cropping system requirements, and environments conditions by a volume of production [25]. Besides individual environment QTL analysis from genome-wide scan in this study (Fig S2), QTL analyses across all individual environments (Fig S3), and within mega-environments were also conducted, which minimized environment effects within MEs (Table S4). This also increased the accuracy to identify a potential major QTL under mega environment and they are very important for local adaptation.

### Dissection of QTL by environment, epistasis and additive-by-environment interactions

Some QTL were very significant for the total LOD score but not for the additive effect LOD scores LOD(A) (Table 1 and 2, Table S4). For example, *Qhi.tamu.4D.29* had LOD(A) of 4.6 among total LOD of 16.9 and the total explained phenotypic variations by the QTL additive effect was 2.5% compared with 13.0% explained by the additive-by-environment. The total additive effect for harvest index was only increased by 0.2%. On the other hand, when analyzed within ME1 including 14EP4, 17BI, 17 CVI, and 17DMS, all irrigated or with high rainfall in that year, the same QTL had LOD(A) of 13.3 from total LOD of 14.1, and variation explained by additive effects was 15.2% from total 19.0% and increased harvest index by 0.6% (Table S4). The same QTL had additive effect that increased harvest index by 1.1% at 17CVI. This is the advantage of dissecting the additive effects from the additive-by-environment interactions to identify the major QTL with higher additive effects but less additive-by-environment interactions. Among the 75 epistasis and epistasis-by-environment interactions that had LOD >= 10 or the interaction effects increased the traits more than those of most major QTL (10.0 g m^−2^ for yield and biomass yield, 0.4 g for thousand kernel weight, 0.4 for kernels spike^−1^, 10 for spikes m^−2^, 0.5% for harvest index, 15 g m^−2^ for biomass yield), only six out of 87 significant QTL involved with the epistasis-by-environment interactions (Table 1 and 2, and Table S5). They were *Qyld.tamu.1B.376, Qtkw.tamu.2D.531, Qhi.tamu.4D.29, Qtkw.tamu.4D.409, Qyld.tamu.6A.12,* and *Qkps.tamu.7B.19,* which can be a warning for breeding selection. Since breeders can only fix the additive effects by selection, through these analyses, breeders can have a better idea for what QTL are worthy of consideration for selection in breeding practice.

### Conclusion

In this study, the wheat 90K Infinium iSelect SNP array and whole genome ddRADseq were used in the construction of high-saturated genetic map for QTL mapping associated with yield and yield components collected from 11 environments across five years and five locations across Texas and New Mexico in the US Southern High Plains. QTL were analyzed using single trait with single environment, single trait across multiple environments, and single trait within and across mega-environments in which lines performed similarly. In addition to additive effects, the interactions of additive-by-environment, epistasis and epistasis-by-environment were dissected. Among 87 significant QTL for nine traits, 36 consistent QTL were identified with presence in at least two above-mentioned analyses and ten pleiotropic QTL were found associated with more than one trait. The eight consistent and pleiotropic QTL were located at 411.7 Mb on chromosome 1A, at 20.6, 26.0, 445.5, 455.3 and 455.8 Mb on chromosome 4D, at 19.6 Mb on 6D and at 64.3 Mb on 7D. They increased dry biomass by 12.1 g m^−2^, harvest index by 0.6%, thousand kernel weight by 0.9 g with favorable alleles from TAM 112 and increased biomass grain yield by 16.3 g m^−2^, kernels spike^−1^ by 0.7, and yield by 20.3 g m^−2^ with favorable alleles from TAM 111. Only six of 75 epistasis-by-environment interactions were involved with the major QTL. Major QTL with larger additive effects and less interaction effects were identified

## Acknowledgments

We thank the technical support from Lisa Garza, Hangjin Yu, Cody Shachter, Srirama Reddy and Chor Tee Tan from the wheat genetic program and graduate students Silvano Assanga and Bharath Reddy from Department of Soil and Crop Science, TAMU.

## Support information

**S1a Fig. Boxplot Analysis of yield and yield component traits.** (DOCX)

**S1b Fig. Dendrogram and biplot Analysis of yield and yield component traits to classify mega-environments for each trait.** (DOCX)

**S2 Fig. LOD profile and additive effects of QTLs detected in each of the 11 environments.** (DOCX)

**S3 Fig. Whole genome significance LOD profiles of quantitative trait loci for yield and its components.** (DOCX)

**S4 Fig. Whole genome significance profiles of epistasis at LOD > 10 for yield and its components.** (DOCX)

**S1 Table. The combined ANOVA, heritability, and mean performance for all traits across environments.** (SLSX)

**S2 Table. Genetic correlation among grain yield and other traits based on overall means and means measured in individual environments.** (SLSX)

**S3 Table. Mapped SNPs on 21 chromosomes, their genetic and physical length.** (SLSX)

**S4 Table. Significant SNPs for yield and yield components in all analysis based on 1000 permutation threshold.** (SLSX)

**S5 Table. Epistases and epistasis by environment interactions for yield and yield components in all analysis.** (SLSX)

## Author Contributions

Conceptualization: Jackie C. Rudd, Amir M. H. Ibrahim, Qingwu Xue, Shuyu Liu.

Data curation: Yan Yang, Smit Dhakal, Chenggen Chu, Shichen Wang, Qingwu Xue, Jackie C. Rudd, Amir M.H. Ibrahim, Kirk Jessup, Jason Baker, Maria Pilar Fuentealba, Ravindra Devkota, Shannon Baker, Charlie D. Johnson, Richard Metz, and Shuyu Liu

Formal analysis: Yan Yang, Smit Dhakal, Chenggen Chu, Shichen Wang, Shuyu Liu.

Funding acquisition: Jackie C. Rudd, Qingwu Xue, Amir M.H. Ibrahim, Yan Yang, Smit Dhakal, Chenggen Chu, Shichen Wang, Shuyu Liu.

Investigation: Smit Dhakal, Maria Pilar Fuentealba, Qingwu Xue, Kirk Jessup, Jackie C. Rudd, Shuyu Liu.

Methodology: Yan Yang, Smit Dhakal, Chenggen Chu, Shichen Wang, Qingwu Xue, Shuyu Liu.

Project administration: Jackie C. Rudd, Qingwu Xue, Shuyu Liu.

Resources: Jackie C. Rudd, Qingwu Xue, Amir M.H. Ibrahim, Charlie D. Johnson, Shuyu Liu.

Software: Yan Yang, Smit Dhakal, Chenggen Chu, Shichen Wang, Shuyu Liu.

Supervision: Shuyu Liu.

Validation: Yan Yang, Smit Dhakal, Shuyu Liu.

Visualization: Yan Yang, Smit Dhakal, Shuyu Liu.

Writing original draft: Yan Yang, Shuyu Liu.

Writing review & editing: Yan Yang, Smit Dhakal, Chenggen Chu, Shichen Wang, Qingwu Xue, Jackie C. Rudd, Amir M.H. Ibrahim, Kirk Jessup, Jason Baker, Maria Pilar Fuentealba, Ravindra Devkota, Shannon Baker, Charlie D. Johnson, Richard Metz, and Shuyu Liu.

